# A putative novel role for *Eip74EF* in male reproduction in promoting sperm elongation at the cost of male fecundity

**DOI:** 10.1101/752451

**Authors:** Sharif Chebbo, Sarah Josway, John M. Belote, Mollie K. Manier

## Abstract

Spermatozoa are the most morphologically variable cell type, yet little is known about genes controlling natural variation in sperm shape. *Drosophila* fruit flies have the longest sperm known, which are evolving under postcopulatory sexual selection, driven by sperm competition and cryptic female choice. Long sperm outcompete short sperm but primarily when females have a long seminal receptacle (SR), the primary sperm storage organ. Thus, selection on sperm length is mediated by SR length, and the two traits are coevolving across the *Drosophila* lineage, driven by a genetic correlation and fitness advantage of long sperm and long SR genotypes in both males and females. *Ecdysone induced protein 74EF* (*Eip74EF*) is expressed during post-meiotic stages of spermatogenesis, when spermatid elongation occurs, and we found that it is rapidly evolving under positive selection in *Drosophila*. Hypomorphic knockout of the *E74A* isoform leads to shorter sperm but does not affect SR length, suggesting that E74A may be involved in promoting spermatid elongation but is not a genetic driver of male-female coevolution. We also found that *E74A* knockout has opposing effects on fecundity in males and females, with an increase in fecundity for males but a decrease in females, consistent with its documented role in oocyte maturation. Our results suggest a novel function of *Eip74EF* in spermatogenesis and demonstrates that this gene influences both male and female reproductive success. We speculate on possible roles for E74A in spermatogenesis and male reproductive success.

**RESEARCH HIGHLIGHTS:** *Eip74EF* promotes oocyte maturation in *Drosophila*. We found evidence that it also promotes sperm elongation in males, but at a cost to male fecundity. Mutant males have shorter sperm but have higher reproductive success, while females have reduced fecundity.

## 1. INTRODUCTION

Spermatozoa are arguably the most morphologically variable cell type among metazoans, exhibiting dramatic diversity in size and shape to encompass all possible permutations of presence or absence of a head, midpiece, or flagellum (Pitnick et al., 2009). With this level of morphological variation, it is not surprising that sperm can evolve rapidly on short time scales (Pitnick et al., 2003; Manier and Palumbi, 2008; Lüpold et al., 2011; Stewart et al., 2016), with morphological differences emerging even over the course of invasive expansion into new environments (Kahrl and Cox, 2017). Sperm morphology can influence swimming speed (Fitzpatrick et al., 2009; Lüpold et al., 2009; Firman and Simmons, 2010; Blengini et al., 2014) and fertilization success (Bennison et al., 2014), and its rapid evolution has been credited to postcopulatory sexual selection via sperm competition (Parker, 1970; Snook, 2005; Pitnick et al., 2009; Godwin et al., 2017) and cryptic female choice (Eberhard, William G., 1986; Firman et al., 2017).

Within the *Drosophila* lineage, sperm length exhibits among the greatest range of phenotypic variation within a clade, spanning over two orders of magnitude from 320 μm in *D. persimilis* to 58,290 μm in *D. bifurca* (Pitnick et al., 1995). Moreover, in the obscura group, males produce heteromorphic sperm, meaning that they come in two (Snook, 1997) or three (Alpern et al., 2019) different sizes within an ejaculate. Sperm length evolution in *Drosophila* is mediated by both length of the female’s primary sperm storage organ, the seminal receptacle or SR (Miller and Pitnick, 2002), a genetic correlation with SR length (Lüpold et al., 2016), and moderate fitness advantage of a long sperm genotype in both males and females (Zajitschek et al., 2019). As a result of this tight interaction between sperm length and SR length, the two traits are coevolving both across species (Pitnick et al., 1999) and among populations undergoing incipient speciation (Pitnick et al., 2003).

Little is known about the genes that control natural variation in sperm length, but we expect them to be rapidly evolving. In a screen for *Drosophila* genes showing signatures of rapid adaptive evolution under positive selection, we identified *Ecdysone-induced protein 74EF* (*Eip74EF*). This gene also has increased expression during post-meiotic stages of spermatogenesis (Vibranovski et al., 2009), when spermatid elongation occurs, though it has no known function in spermatogenesis. Neither isoform E74A nor E74B is expressed in the sperm proteome (Dorus et al., 2006), suggesting that *Eip74EF* is more likely to be involved in regulation of spermatogenesis rather than encoding a structural component of mature sperm. *Eip74EF* is an E26 transformation specific (ETS) domain transcription factor, best understood in its role as an “early response” gene in the ecdysone signaling pathway. Ecdysone regulates many stages of development, including embryogenesis, larval molting, pupation, and metamorphosis (Baehrecke, 1996; Yamanaka et al., 2013) and is also important in early ovary development and virtually all stages of oogenesis (Swevers, 2019). The isoform E74A is specifically required for maintenance and proliferation of germline stem cells during oogenesis (Ables and Drummond-Barbosa, 2010). In spermatogenesis, the ecdysone pathway is critical for the maintenance and proliferation of germline and cyst stem cells (Li et al., 2014; Qian et al., 2014), but it has no known role at later stages.

Here, we explored the role of E74A in sperm length. Because sperm length and SR length are genetically correlated (Lüpold et al., 2016) and coevolving (Pitnick et al., 1999, 2003), we also measured SR length in E74A hypomorphic mutant females to look for pleiotropic effects on both sperm length and SR length. We found that E74A mutant males had shorter sperm, so we also examined mating behavior and sperm competitive success in mutant males and fecundity in both males and females.

## 2. MATERIALS AND METHODS

### 2.1 Molecular evolution

We estimated the rate of positive selection on *Eip74EF* as a measure of adaptive molecular divergence using DNA sequences from 22 *Drosophila* species downloaded from NCBI. The codeml program in PAML v4.6 (Yang, 1997, 2007) compares a null model of neutral divergence (M8a) with a model of adaptive divergence under positive selection (M8), based on the ratio *dN*/*dS*, which compares the rate of non-synonymous substitutions (*dN*) with the rate of synonymous substitutions *(dS). dN/dS* = ω, and generally *ω* < 1 points to purifying selection, *ω* = 1 indicates neutrality, and *ω* > 1 suggests positive selection. However, *ω* is rarely greater than 1, since multiple sites across multiple lineages are unlikely to be under positive selection for an extended period. Thus, ω estimates are conservative indicators of positive selection on codons. The critical values for this test are 2.71 at 5% and 5.41 at 1%, with a null distribution of point mass 0 and *χ*^2^_1_ (Self and Liang, 1987).

### 2.2 Stocks and crosses

*Eip74EF* has only one major protein isoform expressed in adults, E74A (Liu et al., 2012). For this study, we used a hypomorphic mutant of *Eip74EF* obtained from the Bloomington Drosophila Stock Center (stock #12619) that has partial reduction of expression of E74A while retaining viability (Liu et al., 2012). The mutant allele (*E74A^BG01805^*) contains a *p*(*GT1*) transposon insertion in the sixth intron, within a *w^1118^* background. Experimental animals were reared from breeding vials containing 10 virgin females and 20 virgin males that were transferred to new sugar-yeast-agar food vials daily for four days, after which adults were discarded. Control animals were reared from breeding vials containing 10 *E74A^BG01805^* females and 20 *w^1118^* males. Progeny were collected within 8-12 hours of eclosion using light CO2 anesthesia into single-sex vials at densities of 10 females or 20 males per vial. For both experimental and control treatments, progeny from two replicate sets of breeding vials out of four potential replicates per treatment were phenotyped for all experiments.

For reproductive success experiments, competitor males were derived from a Canton-S stock expressing a 3xP3-GFP construct in eye. These Canton-S Protamine-GFP lines were generated using the piggybac germline transformation system (Handler and Harrell, 1999). The GFP-tagged Protamine-B construct was cut out of the *P*-element plasmid construct pW8/ProtB-GFP1 (Manier et al., 2010) with *Kpn*I and *Bam*HI and subcloned into the pBS/2xAsc vector, a modified pBlueScript-KS+ plasmid (Stratagene, La Jolla, CA) in which two *Asc*I sites, flanking the multiple cloning site, had been created by site-directed mutagenesis. The insert was then cut out with *Asc*I and ligated into the single *Asc*I site of the piggybac vector p3xP3/EGFPaf (Horn and Wimmer, 2000) to yield p3xP3-EGFP/ProtB-GFP. This DNA was mixed with the helper plasmid pBΔSac (Handler and Harrell, 1999) and injected into preblastoderm embyos from a Canton-S wild-type strain (generously provided by Mariana Wolfner, Cornell University). Transformants were identified by green fluorescent pseudopupils and ocelli. Two tagged lines were generated: *Canton-S pBac{3xP3-EGFP, ProtB-EGFP}27* and *Canton-S pBac{3xP3-EGFP, ProtB-EGFP}16B* with the transgenes inserted on chromosome 2 and chromosome 3, respectively. Although this stock (hereafter “Canton-S”) contained a transgene, the fluorescent eye and sperm markers were not specifically utilized in this series of experiments.

### 2.3 Sperm measurements

To obtain sperm length measurements, seminal vesicles from 5-day-old ether-anesthetized males were dissected into a large droplet of 1X phosphate-buffered saline (PBS) on a gelatin-coated glass slide, using fine forceps. The seminal vesicle was ruptured and dragged several times through the droplet to release and spread motile sperm. The slide was then dried down at 55 °C in a drying oven, fixed in 3:1 methanol:acetic acid, stained with DAPI (Molecular Probes), and mounted under a coverslip in 80% glycerol in PBS and sealed with nail polish. We then visualized sperm using darkfield microscopy at 200X and sperm heads using fluorescence microscopy at 400X on a Nikon Ni-U microscope and captured images with a Zyla Andor monochrome camera. Images were measured using the segmented line tool in ImageJ software (https://imagej.nih.gov/ij/). Up to five sperm and sperm heads were measured for each of up to 8 males across two replicates for both the mutant and control treatments. Sperm measurements that were abnormally short for a given male were assumed to be due to breakage and were excluded.

### 2.4 SR length

Females were aged 5 days and then frozen in their food vials until dissection. Females were then thawed, and thorax length was measured as a proxy for body size using a ruler reticle in a Nikon SMZ-745 stereoscope at maximum magnification. The lower reproductive tract was dissected with fine forceps into a droplet of 1X PBS, and the SR was carefully unraveled using a fine insect pin. The sample was then mounted under a coverslip with clay on the corners to allow compression of the sample into a single plane without overcompression. SRs were immediately visualized at 100X or 200X under phase contrast microscopy, and images were captured using a Canon EOS Rebel T3i camera. For each of two replicates for mutant and control treatments, 10 females were measured using the segmented line tool in ImageJ.

### 2.5 Fitness

We evaluated the fitness consequences of *Eip74EF* knockdown by quantifying male and female fecundity and male competitive fertilization success. To measure fecundity, we housed 20 males or females per replicate individually with two Canton-S females or males, respectively, for seven days, transferring triads (one focal individual and two mates) into new food vials every two days, without anesthesia. We counted adult progeny that eclosed from all food vials. This approach to counting progeny to assess fecundity does not account for effects of *Eip74EF* on egg-to-adult viability, only on the total number of adult progeny that survived to successfully eclose.

We assessed paternity success of mutant and control males by mating a *w^1118^* female first with a Canton-S male (day 0) and giving her a four-hour opportunity to remate with a mutant or control male over the following 4 days (days 1-4). All matings occurred in individual vials and were observed directly, recording copulation duration and day of remating. We scored paternity using eye color, since females, mutant males, and control males carried a recessive white-eye mutation, while Canton-S competitor males had a dominant wild type eye color. Paternity was expressed as P2, or proportion of offspring sired by the second male (here, mutant or control males).

### 2.6 Statistical analysis

All statistical analyses were performed in R v.3.5.3 (Team, 2018). Total sperm length and sperm head length were analyzed using lmer within library lme4, with knockdown treatment as a fixed effect, male identity nested within replicate as a random effect, and researcher identity as a random effect, with restricted maximum likelihood (REML) parameter estimation. SR length was found to be negatively correlated with body size (thorax length) (Pearson’s product-moment correlation, *t*_42_ = −2.514; *P* = 0.0159), so thorax length was included as a random effect in a linear mixed model, with knockdown treatment and replicate as fixed effects, and thorax length as a random effect. SR lengths were all measured by the same individual. Fecundity data were analyzed using lmer with knockdown treatment as a fixed effect and replicate as a random effect, with maximum likelihood parameter estimation, because there was only one random effect. For all analyses implemented in lmer, P-values were estimated using a Type II Wald chi-square test with Anova in the car library. Here, we report knockdown treatment effect sizes as the *t*-value from the mixed model regression, along with the χ^2^ and p-value from the Wald’s test.

From the sperm competition experiment, we also obtained remating rate (proportion of females that remated), copulation duration, and day of remating. Remating rate was analyzed using a chi-square test in two different ways: with replicates as categories (N = 4) and with knockdown treatments as categories (N = 2), both yielding the same result. Copulation duration was analyzed using lmer with knockdown treatment as a fixed effect and replicate as a random effect, with maximum likelihood parameter estimation. We report results as for other lmer analyses described above. For paternity success, we used logistic regression with a logit link function and binomial error distribution (after ensuring no overdispersion in the data), implemented using glm.

## 3. RESULTS

*Eip74EF* is rapidly evolving under positive selection, with an M8a model estimate of −20882.5, M8 of −20875.6, and a *χ*^2^ of 13.80, well above the 1% critical threshold of 5.41. Knockdown of *Eip74EF* resulted in shorter sperm (*t* = −5.341, *χ*^2^_1_ = 28.53, *P* < 0.0001; Figure 1a) but had no effect on sperm head length (*t* = −1.48, *χ*^2^_1_ = 2.191, *P* = 0.139; Figure 1b) nor female SR length (*t* = 1.132, *χ*^2^_1_ = 1.281, *P* = 0.258; Figure 1c). Disruption of *E74A* led to an increase in male fecundity (*t* = 3.529, *χ*^2^_1_ = 12.45, *P* = 0.0004; Figure 2a) but a decrease in female fecundity (*t* = −10.17, *χ*^2^_1_ = 103.43, *P* < 0.0001; Figure 2b). In other words, female fecundity was adversely affected while male fecundity was positively affected. In the sperm competition experiment, female remating rate (replicates as groups: *χ*^2^_3_ = 3.143, *P* = 0.370; treatment as groups: *χ*^2^_1_ = 0.762, *P* = 0.383; Figure 3a), day of remating (*t* = 0.336, *χ*^2^_1_ = 0.113, *P* = 0.737; Figure 3b), copulation duration (*t* = 0.109, *χ*^2^_1_ = 0.012, *P* = 0.913; Figure 3c), and paternity success (P_2_; *z* = 0.163, *P* = 0.871; Figure 3d) were all unaffected by E74A knockout.

**FIGURE 1.**
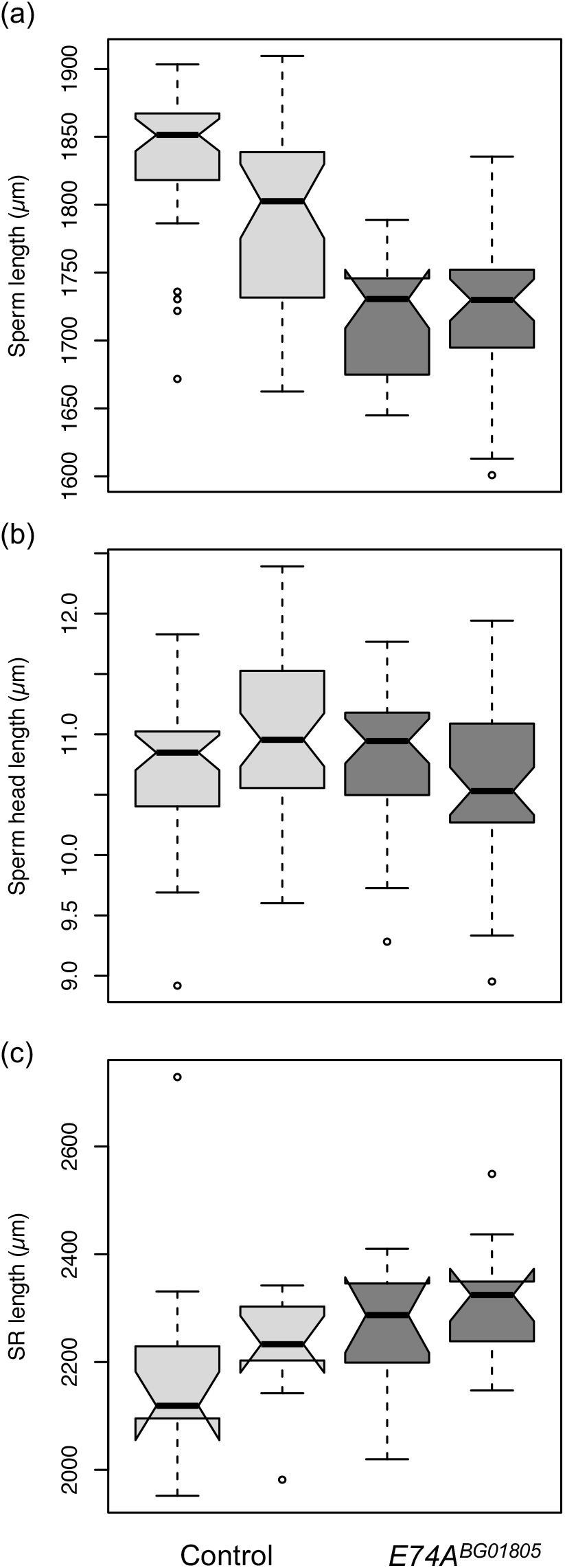
*E74A^BG01805^* males had significantly shorter sperm (*P* < 0.0001) (a) with no difference in sperm head length (*P* = 0.139) (b), and *E74A^BG01805^* females had no change in SR length (*P* = 0.258) (c) relative to controls. Two replicates are shown per treatment, with the mutant in dark gray and control in light gray.

**FIGURE 2.**
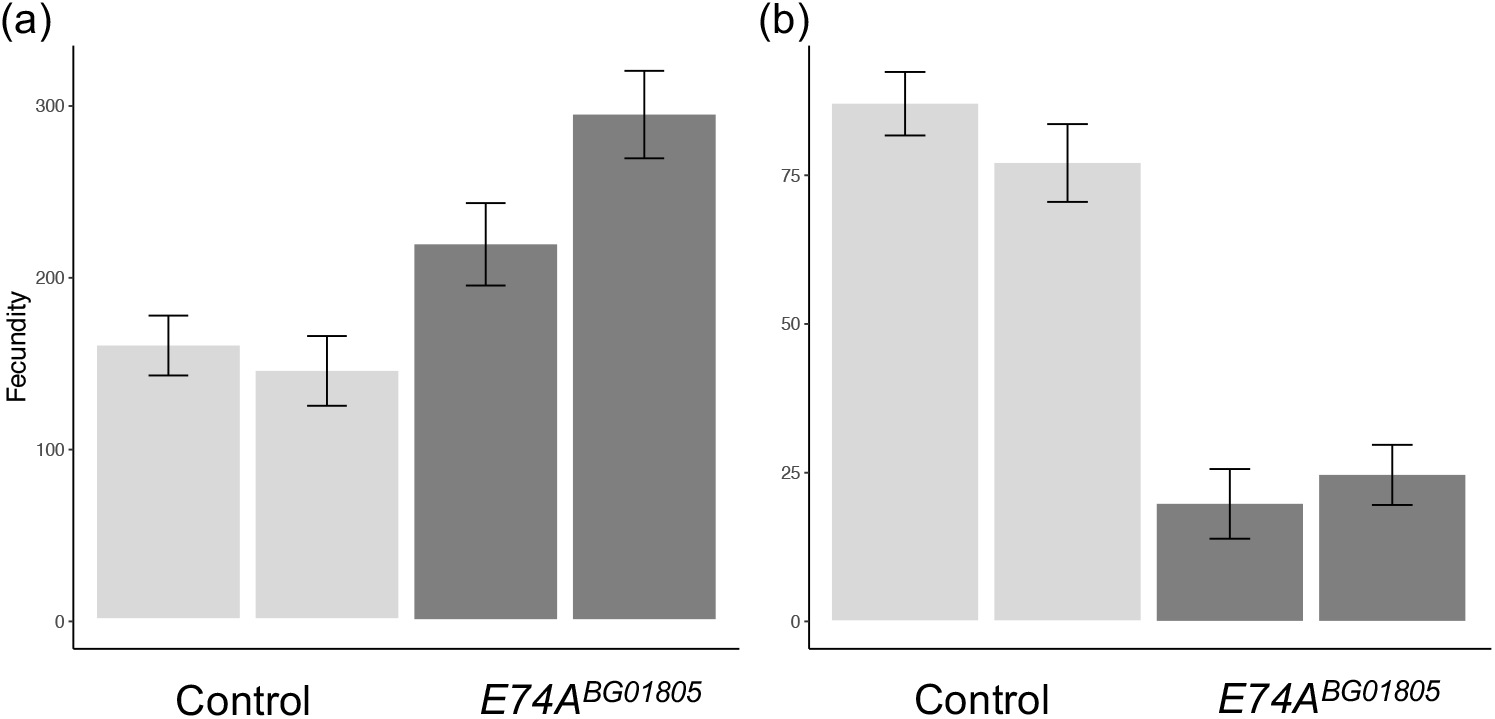
*E74A^BG01805^* males had higher fecundity (P = 0.0004) (a), while *E74A^BG01805^* females had lower fecundity (P < 0.0001) (b). Two replicates are shown per treatment, with the mutant in dark gray and control in light gray.

**FIGURE 3.**
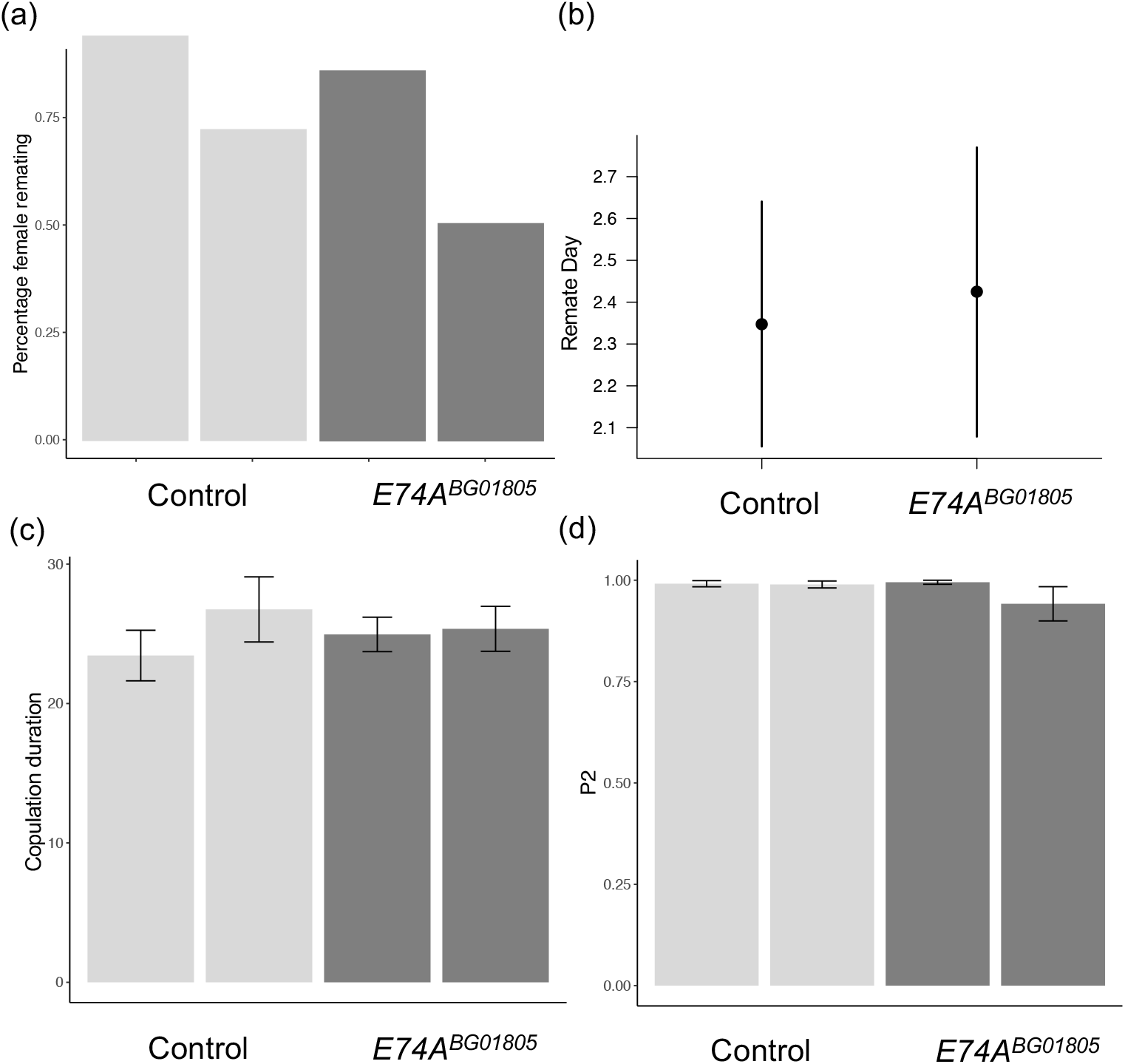
The *E74A^BG01805^* mutation had no effect on a male’s ability to influence female remating rate (*P* = 0.323) (a), female latency to remate in days (*P* = 0.737), illustrated as effect sizes of the statistical model (b), copulation duration in minutes (*P* = 0.913) (c), and P2 (*P* = 0.871) (d). Two replicates are shown per treatment, with the mutant in dark gray and control in light gray.

## 4. DISCUSSION

We found that *Eip74EF* is evolving rapidly under positive selection, a surprising result given that its roles in molting and metamorphosis, which are both highly conserved processes, would suggest that it should be under strong purifying selection. However, we found evidence that E74A plays a novel role during sperm morphogenesis, in the positive regulation of sperm length, a rapidly evolving trait. The decrease in sperm length in *E74A* mutants is attributable primarily to change in sperm flagellum length, since head length was unaffected. At the same time, mutant males had increased fecundity. It should be noted that we have only included one genetic experiment here, and our sperm length results may be due to other genetic factors linked to *Eip74EF* rather than this gene itself. More experimentation is needed to further confirm the effect of E74A on sperm length including, but not limited to, examining sperm length in a knock-in mutant with expression under control of a heat-shock promoter (Fletcher et al., 1997).

We know little about E74A’s role in testis. Within the ecdysone pathway more generally, it is one of several genes that are activated when the steroid hormone 20-hydroxyecdysone (ecdysone) binds to a heteromeric receptor in the nuclear envelope comprised of ecdysone receptor (EcR) and ultraspiracle (USP) (Yamanaka et al., 2013). EcR is also a transcription factor that directly initiates expression of a number of early response genes, including *Eip74EF* (Uyehara and McKay, 2019). In the male germline, EcR is required for growth and proliferation of both germline stem cells and cyst stem cells, which co-differentiate throughout spermatogenesis (Li et al., 2014; Qian et al., 2014). Further indirect evidence for E74A function in spermatogenesis comes from defects found in *belle* (*bel*) mutants. *Bel* is a DEAD box RNA helicase that regulates translation of *E74A* transcripts in salivary glands during metamorphosis (Ihry et al., 2012). Although it is unknown if *bel* similarly regulates *E74A* translation in testis, hypomorphic mutants of *bel* have spermatogenic defects in chromosome segregation and cytokinesis at the round spermatid stage, approximately when spermatid elongation begins (Johnstone et al. 2005).

We can only speculate about E74A’s possible function during spermatogenesis. We do know that it is a transcription factor that regulates downstream genes in the ecdysone signaling pathway. Ecdysone signaling generally tends to promote the maintenance of developmental stages or to trigger a switch between stages (Gancz and Gilboa, 2013; Li et al., 2014; Yatsenko and Shcherbata, 2018; Leiblich et al., 2019; Swevers, 2019), with parallel functions in larval/ pupal development and gametogenesis (Ables et al., 2016). Because germline development is regulated largely by non-autonomous paracrine signaling from supporting somatic cells (Spradling et al., 2011; Zoller and Schulz, 2012; Li et al., 2014; Qian et al., 2014; Ables and Drummond-Barbosa, 2017), E74A is likely to be expressed in somatic cyst cells. Indeed, ecdysone signaling is detectable in nurse cells and follicle cells throughout oogenesis (Carney and Bender, 2000) and regulates multiple stages from development of the ovarian niche (Gancz and Gilboa, 2013; Yatsenko and Shcherbata, 2018) to ovulation (Knapp and Sun, 2017; Swevers, 2019). In the ovary, ecdysone signaling is orchestrated on a fine spatio-temporal scale throughout oogenesis (Swevers 2019). In spermatogenesis, E74A is also likely to be tightly regulated at very specific timepoints, especially since expression in the testis overall is very low (Brown et al., 2014; Li et al., 2014). Less is known about the role of ecdysone signaling in morphogenesis, though it is required for morphogenesis and function of Malpighian tubules (Gautam et al., 2015). Perhaps most relevant to flagellar elongation, an early study on *Drosophila* Kc cells in culture found that application of ecdysone induced elongation and motility in cells mediated by the formation of filopodia through tubulin assembly (Berger et al., 1980). This result suggests that ecdysone could also mediate flagellar assembly during spermatogenesis.

On the face of it, it is curious that knockdown of *Eip74EF* has opposing consequences in males and females, with increased fecundity in males but decreased fecundity in females. The result in females is more easily explained by the role that *Eip74EF* plays in oogenesis; without it, germline stem cells arrest development and degenerate. In males, *Eip74EF* seems to promote longer sperm but at a cost to male fecundity. There are a number of events between mating and oviposition during which fecundity could ultimately be affected, including sperm transfer, sperm storage, sperm retention in storage, ovulation, sperm release from storage, fertilization, oviposition, and offspring viability to adulthood. It is possible that there is a trade-off in *Eip74EF* mutants between producing longer sperm and producing more sperm, such that because mutants produce shorter sperm, they are now able to produce more sperm (Pitnick and Markow, 1994; Immler et al., 2011). This hypothesis is unlikely to explain the whole story, however, since fecundity in mutant males increased by approximately 50%, while sperm length only decreased by around 5%. Further experiments should examine the rate of spermatogenesis in testes of E74A mutant males.

It is also possible that fewer mutant sperm are able to enter storage. Decreased storage may be a function of sperm storage organs being able to store more shorter sperm. An alternative and non-mutually exclusive hypothesis is that E74A regulates seminal fluid composition and/or transfer. Seminal fluid proteins produced in the accessory glands can have profound effects on many aspects of female reproduction, behavior, and physiology (Avila et al. 2011), including sperm storage after mating (Neubaum & Wolfner 1999; Bloch Qazi & Wolfner 2003; Avila & Wolfner 2009). Accessory gland development and function require ecdysone receptor (EcR) (Herndon et a. 1997; Sharma et al. 2017; Leiblich et al 2019), which directly upregulates E74A in salivary glands and wing disc (Burtis et al., 1990; Thummel et al., 1990; Uyehara and McKay, 2019). It is therefore possible that E74A is also involved in accessory gland function and production of seminal fluid proteins independent of a possible role in morphogenesis during spermatogenesis.

Despite having shorter sperm, *E74A* hypomorphic mutants were not less competitive during sperm competition. Previous work found that the long-sperm advantage only occurs when mating with females with long SRs. We did not measure the SR length of females used in the paternity study, nor did we measure sperm length in the standard male competitors. It is possible that the combinations of male sperm and female SR phenotypes were not conducive to observing a fitness effect of E74A under sperm competition. More controlled experiments in which knockout males competed against males of known sperm length within females of known SR length would more clearly demonstrate a role for E74A in sperm competition.

What is driving the rapid evolution of *Eip74EF*? On an evolutionary level, it may reflect selection on sperm length, mediated by length of the SR (Miller and Pitnick, 2002) and a genetic correlation between the two traits (Lüpold et al., 2016). However, most reproductive proteins that are rapidly evolving are ones that interact directly with female-derived molecules, such as gamete-recognition proteins that mediate fertilization and seminal fluid proteins that modify and are modified by molecules in the female reproductive tract (Wilburn and Swanson, 2016). Here, we demonstrate rapid evolution of a protein that has no known direct interactions with female-derived molecules, nor does it seem to affect length of the SR, which does directly interact with sperm length. Dorus et al. (2006) found that many sperm proteome proteins are well-conserved, suggesting that structural proteins that form the building blocks of spermatozoa are evolving slowly, while the molecular builders that regulate sperm morphogenesis evolve more rapidly. Thus, the molecular secret to rapid evolution and the vast morphological diversity of sperm may lie not in its final structure but in its development, harkening back to the old adage of evo-devo: we cannot understand the evolution of morphological diversity until we understand the evolution of development.

## Abbreviations

*Eip74EF*: Ecdysone induced protein 74EF
GFP: green fluorescent protein
NCBI: National Center for Biotechnology Information
PAML: Phylogenetic Analysis by Maximum Likelihood
PBS: phosphate buffered saline
REML: restricted maximum likelihood
SR: seminal receptacle

## DATA AVAILABILITY STATEMENT

The data that support the findings of this study are openly available in Dryad at https://doi.org/10.5061/dryad.k0p2ngf4k.

## ACKNOWLEDGEMENTS

Experimental assistance was provided by Michael DeNieu, Gary Hovsepian, Paul Kwon, Isvita Marfatia, and Ponmali Photavath. This work was funded by NSF DEB-1257859 to MKM and JMB, and the George Washington University Harlan Undergraduate Research Fellowship to SC. The authors declare no conflicts of interest.

